# Impact of acetolactate synthase inactivation on 1,3-propanediol fermentation by *Klebsiella pneumoniae*

**DOI:** 10.1101/365387

**Authors:** Sheng Zhou, Youhua Huang, Xinliang Mao, Lili Li, Chuanyu Guo, Yongli Gao, Qiwei Qin

## Abstract

1,3-Propanediol (1,3-PDO) is an important compound that is mainly used in industry for polymer production. Fermentation of 1,3-PDO from glycerol by marine *Klebsiella pneumoniae* is accompanied by formation of 2,3-butanediol (2,3-BDO) as one of the main byproduct. The first step in the formation of 2,3-BDO from pyruvate is catalyzed by acetolactate synthase (ALS), an enzyme that competes with 1,3-PDO oxidoreductase for the cofactor NADH. This study aimed to analyze the impact of engineering the 2,3-BDO formation pathway via inactivation of ALS on 1,3-PDO fermentation by marine *K. pneumoniae* HSL4. An ALS mutant was generated using Red recombinase assisted gene replacement. The ALS specific activities of *K. pneumoniae* ΔALS were notably lower than that of the wild-type strain. Fed-batch fermentation of the mutant strain resulted in a 1,3-PDO concentration, productivity and conversion of 72.04 g L^−1^, 2.25 g L^−1^ h^−1^, and 0.41 g g^−1^, a slightly increase compared with the parent strain. Moreover, inactivation of ALS decreased *meso*-2,3-BDO formation to trace amounts, significantly increased 2S,3S-BDO and lactate production, and a pronounced redistribution of intracellular metabolic flux was apparent.

## 1 Introduction

1,3-Propanediol (1,3-PDO) is an important industrial chemical intermediate that is used in the synthesis polymers [1, 2]. 1,3-PDO can be produced by both chemical synthesis and microbial fermentation. While chemical methods are troubled by poor selectivity and high energy consumption, microbial fermentation has a number of advantages including mild reaction conditions that are more environmentally favourable, greater sustainability, and easier operation [3, 4]. Numerous microorganisms produce 1,3-PDO including *Klebsiella pneumoniae*, *Citrobacter freundii, Clostridium butyricum*, and *Clostridium acetobutylicum* [5-7]. Of these, K.*pneumoniae* has been most widely explored since it achieves a higher concentration of 1,3-PDO, greater productivity, and a higher rate of conversion using glycerol as the carbon source [8, 9].

In K. *pneumoniae* metabolism, glycerol is dissimilated through coupled reductive and oxidative pathways (Fig. 1). 1,3-PDO is synthesized in the reductive pathway, whereas in the oxidative pathway, pyruvate-derived by-products such as 2,3-butanediol (2,3-BDO), acetate (AC), lactate (LAC), succinate (SUC) and ethanol (ETH) are generated through the channeling of glycerol into glycolysis [10]. Generation of by-products can compete with the biosynthesis of 1,3-PDO for carbon flow and NADH, which can lower the yield of 1,3-PDO. Blocking both the ETH and LAC pathways can reduce the production of by-products and can increase 1,3-PDO [11-13].

**Figure 1.**
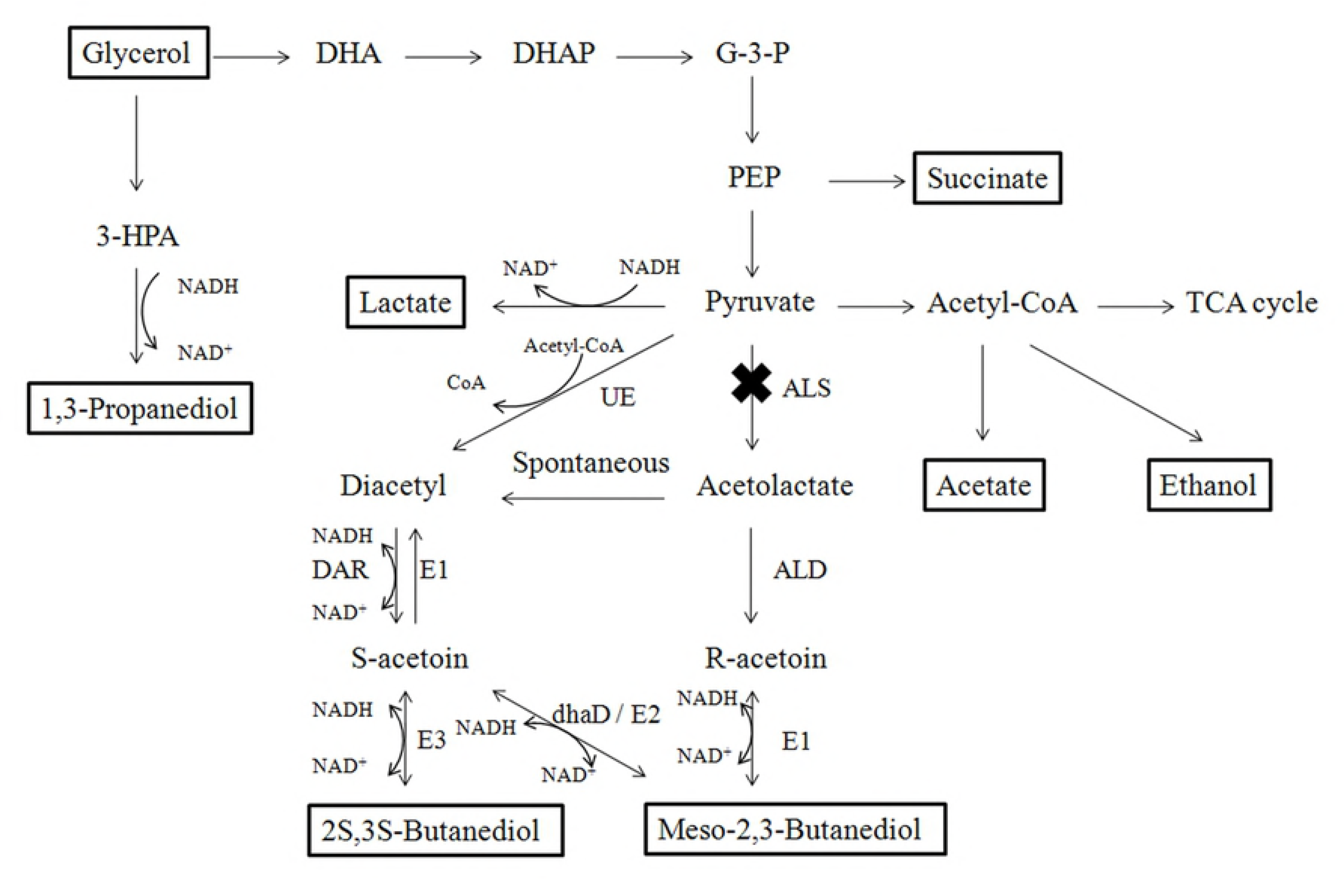
Metabolic pathways of glycerol metabolism and the proposed mechanism for the biosynthesis of 2,3-BDO stereoisomers in K. *pneumoniae* [28, 29]. The glycerol substrate and major product metabolites are boxed. UE: Unknown enzyme; ALS: acetolactate synthase (mutated in this study); ALD: α-acetolactate decarboxylase; DAR: diacetyl reductase; E1: meso-2,3-BDO dehydrogenase (R-acetoin forming); E2: *meso*-2,3-BDO dehydrogenase (S-acetoin forming); E3: (S,S)-2,3-BDO dehydrogenase; dhaD: Glycerol dehydrogenase.

2,3-BDO is the main byproduct generated alongside 1,3-PDO in the oxidative catabolism of glycerol, and both compounds share a similar high boiling point and water solubility. Therefore, 2,3-BDO not only competes for carbon flow, but also complicates 1,3-PDO recovery and purification [14]. Minimizing 2,3-BDO production would therefore improve the yield of 1,3-PDO and simplify the downstream processing [15, 16].

2,3-BDO is a metabolite derived from pyruvate (Fig. 1). In the *Enterobacteriaceae* family, pyruvate from glycolysis can be converted into either LAC in an NADH-dependent reaction catalyzed by L-/D-LAC dehydrogenase (LDH) or, after decarboxylation, into acetolactate in a reaction catalyzed by acetolactate synthase (ALS), encoded by the budB gene. Acetolactate is mostly produced when NADH availability is low, and under aerobic conditions, it can be converted to acetoin by acetolactate decarboxylase (ALD), encoded by the budA gene. If oxygen is present, acetolactate can undergo spontaneous decarboxylation to produce diacetyl, and diacetyl reductase (DAR; also known as acetoin dehydrogenase) converts diacetyl to acetoin. Finally, butanediol dehydrogenase (BDH), encoded by the budC gene, also known as acetoin reductase, reduces acetoin to 2,3-BDO [17].

In this study, a 2,3-BDO pathway-deficient mutant of *K. pneumoniae* was constructed by knocking out the budB gene to generate a variant strain lacking ALS. The physiological and fermentation properties of the mutant strain, including the distribution and yield of various metabolites and products, were subsequently investigated. In particular, the influence that blocking the 2,3-BDO pathway has on glycerol metabolism and 1,3-PDO production was studied.

## 2 Materials and methods

### 2.1 Strains, plasmids and culture media

Strains and plasmids used in this study are listed in Table 1. *Klebsiella pneumoniae* HSL4, used for the biosynthesis of 1,3-PDO in this work, was previously isolated from mangrove sediment samples [9]. Basic culture media contained the following components (L^−1^): 16 g pure glycerol (mass percent > 99%), 1.5 g yeast extract, 4 g (NH_4_)_2_SO_4_, 0.69 g K_2_HPO_4_, 0.25 g KH_2_PO_4_, 0.2 g MgSO_4_·7H_2_O, 1 ml trace element solution, and 1 ml Fe^2+^ solution. The trace element solution contained (L^−1^): 100 mg MgSO_4_·4H_2_O, 70 mg ZnCl_2_, 35 mg Na_2_MoO_4_·2H_2_O, 60 mg H_3_BO_3_, 200 mg CoCl_2_·6H_2_O, 29.28 mg CuSO_4_·5H_2_O, 25 mg NiCl_2_·6H_2_O, and 0.9 ml HCl (37%).

**Table 1.** Bacterial strains and plasmids.

| Strain or plasmid | Relevant genotype and description | Source or reference |
| --- | --- | --- |
| Strains |  |  |
| *K. pneumoniae* HSL4 | Wild-type, CCTCC M 2011075 | [9] |
| *K. pneumoniae*-pDK6-red | *K. pneumoniae* CCTCC M2011075, pDK6-red | [18] |
| *K. pneumoniae* ΔALS | *K. pneumoniae* CCTCC M2011075, ΔALS, aac(3)IV | This work |
| *Escherichia coli* DH5a | Host of plasmid | Lab stock |
| Plasmids |  |  |
| pMD18-T simple | Amp<sup>r</sup>, TA cloning vector, 2692 bp | TaKaRa |
| pMD18-T-ALS | Amp<sup>r</sup>, carries ALS, 4231 bp | This work |
| pMD18-T-ΔALS | Amp<sup>r</sup>, carries ΔALS, 5230 bp | This work |
| pDK6-red | Kan<sup>r</sup>, carries λ-Red genes (gam, bet, exo), 7.1 kb | [18] |
| pDK6 | Kan<sup>r</sup>, lacIQ, tac, 5.1 kb | [19] |
| pKD46 | Amp<sup>r</sup>, λ Red recombinase expression | [20] |
| pIJ773 | Apra<sup>r</sup>, aac(3)IV with FRT sites, 4334 bp | [21] |
| pIJ790 | Cm<sup>r</sup>, encodes λ-Red genes (gam, bet, exo), 6084 bp | [21] |

When necessary, the medium was supplemented with antibiotics at the following final concentrations: 50 μg mL^−1^ kanamycin (kan), 100 μg mL^−1^ ampicillin (amp), 50 μg mL^−1^ apramycin or 25 μg mL^−1^ chloramphenicol.

### 2.2 Construction of mutant strain *K. pneumoniae* ΔALS

The ALS mutant strain K. *pneumoniae* ΔALS was constructed following a method described by Wei et al. [18]. Briefly, the ALS gene of *K. pneumoniae* HSL4 was amplified by PCR using primer pair ALS-F1/ALS-R1. The PCR product was ligated into the pMD18-T simple vector to generate pMD18-T-ALS. A linear portion of DNA with both 40 nt homologous extensions flanking the apramycin resistance gene aac(3)IV was amplified using plasmid pIJ773 as the template with primer pair ALS-F2/ALS-R2. The pMD18-T-ΔALS construct was engineered by replacing part of the ALS gene in plasmid pMD18-T-ALS with the aac(3)IV cassette using the Red/ET system in *Escherichia coli* [20]. The pMD18-T-ΔALS vector was used as a template for the PCR-based preparation of a linear DNA containing the apramycin resistance gene aac(3)IV with 468 bp upstream and 714 bp downstream homologous regions. Finally, the linear DNA product was transformed into K. *pneumoniae*-pDK6-red, which already hosted the pDK6-red plasmid. Homologous recombination between the introduced linear DNA and chromosomal DNA was facilitated by Red recombinase and led to the deletion of ALS (Fig. 2). The ALS-deficient strains were selected on apramycin plates, and primer pairs Apra-F/Apra-R and ALS-F3/ALS-R3 were used for PCR confirmation. Primers used in this study are listed in Table 2.

**Figure 2.**
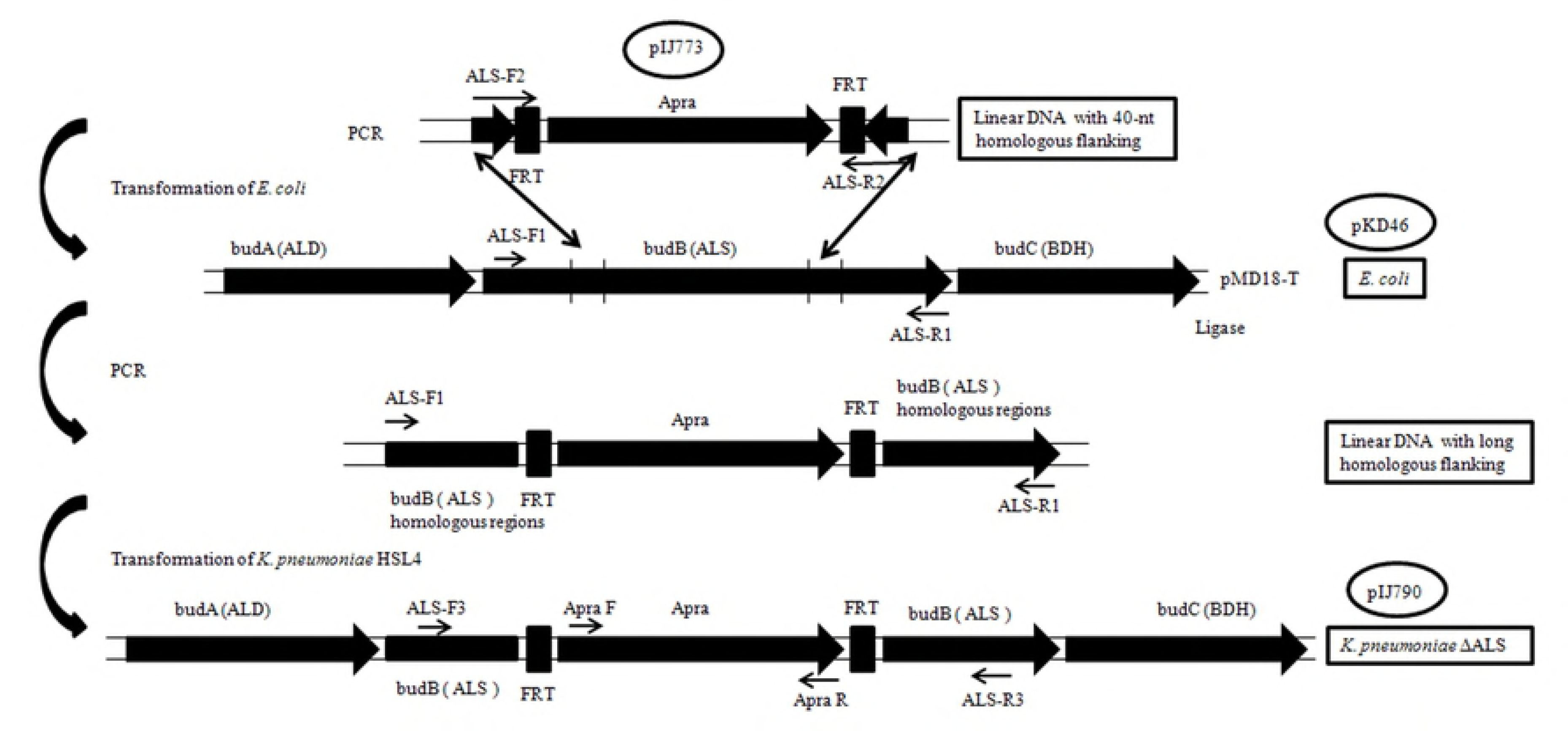
Methodology for ALS gene mutation using Red recombination. BudA (ALD) and budC (BDH) are upstream and downstream of ALS on the K. *pneumoniae* HSL4 chromosome, respectively. ALS-F1/R1, ALS-F2/R2, ALS-F3/R3, and Apra F/R were the primer pairs used.

**Table 2.**
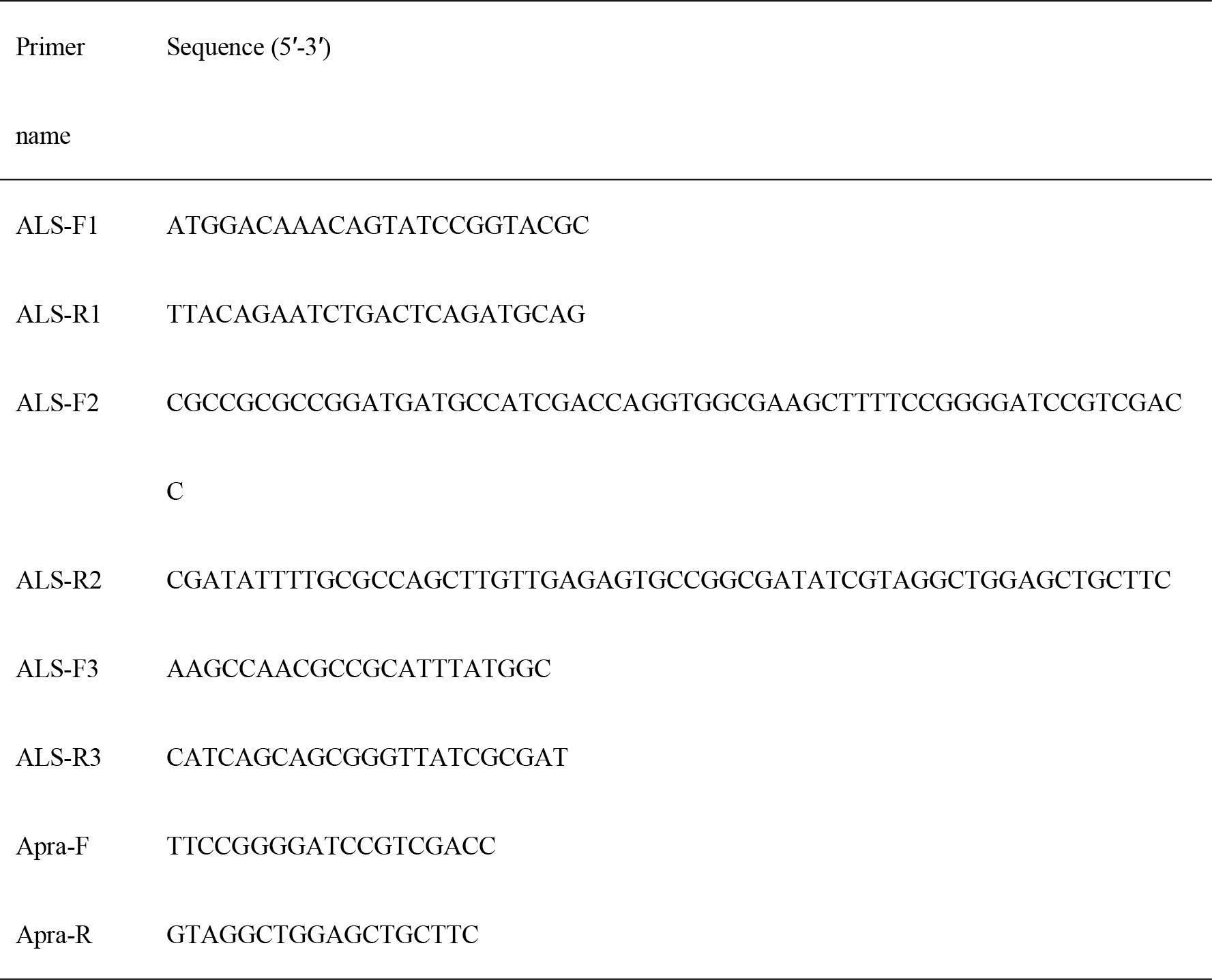
Primers used in this study.

### 2.3 Batch and fed-batch fermentation

For batch fermentation and preparation of seed cultures for fed-batch fermentation, 250 mL flasks containing 100 mL of basic culture medium (glycerol adjusted to 30 g L^−1^) were inoculated with a single colony and incubated at 37°C in a rotatory shaker at 200 rpm for 12 h. For fed-batch fermentation, a 3% (v/v) inoculum was added aseptically to a 5 L fermenter (Zheng Jiang Dong Fang, China) containing 3 L of media. NaOH (8 M) was used to maintain the pH. Aerobic fermentation Hao et al. [22] was carried out at 37°C, pH 6.8, 200 rpm with an air flow of 0.5 vvm (volume per volume per minute). The glycerol concentration was 16–18 g L^−1^ at the beginning of fermentation, and when it dropped below 5 g L^−1^, additional pure glycerol was added to a concentration of 2–8 g L^−1^ and 8–20 g L^−1^ in subsequent fed-batch cultures at 6–16 h and 16–32 h, respectively.

### 2.4 The ALS activity assays

The ALS assay was based on enzymatic conversion of pyruvate to acetoin through acetolactate, and was conducted as described by Yang et al. [23]. Samples were collected and centrifuged at 5000 × *g*, 4°C for 10 min, supernatants were removed, and pellets were stored at 4°C until needed for analysis. Pellets were washed with cold 50 mM acetate buffer pH 6, centrifuged at 5000 × *g*, 4°C for 10 min, resuspended in the same buffer at 4°C, and crude cell extracts (for use in enzyme assays) were obtained by sonication with an ultrasonic cell disrupter. Reaction mixtures contained 20 mM sodium pyruvate, 50 mM acetate buffer pH 6, 1 mM MgCl_2_, and 80 mL of cocarboxylase. Reactions were initiated by the addition of 100 μL of crude extract to 1 mL of reaction buffer and followed for 20 min at 37°C. Reactions were terminated by addition of 0.1 mL of 50% H_2_SO_4_. Mixtures were incubated for an additional 25 min at 37°C to allow acid hydrolysis of acetolactate to acetoin, and acetoin was measured using the Voges–Proskauer test. Briefly, a 0.2 mL aliquot of the aforementioned mixture was transferred to a tube with 0.8 mL of 0.45 M NaOH, and 1 mL of α-naphtholf (5% in 2.5 M NaOH) and 0.5% creatine (1:1) was added. Reactions were incubated at room temperature for 30 min with slightly shaking, and the absorbance at 535 nm was measured. Calibration with pure acetoin showed a correlation of 1 OD_535_ unit to 0.15 mM acetoin. One specific unit of ALS activity corresponds to the formation of 1 mmol of acetoin per mg of protein per minute.

### 2.5 Biomass and fermentation broth analysis

Cell density was monitored using a spectrophotometer at 600 nm (OD600nm). Glycerol, 1,3-PDO, 2,3-BDO, LAC, SUC, AC and ETH were determined using a Shimadzu 20AT HPLC system equipped with a 300 × 7.8mm^2^ Aminex HPX-87H column (Bio-Rad, USA) running at 0.8 ml min^−1^ after adding 5 mM H_2_SO_4_.

## 3 Results

### 3.1 Inactivation of the ALS gene in K. *pneumoniae* HSL4

A Red recombination approach was used to knock-out the ALS gene in K. *pneumoniae* HSL4 (Fig. 2). A positive recombinant was selected from the apramycin plate, confirmed by PCR and named *K. pneumoniae* ΔALS.

### 3.2 Effect of ALS inactivation on cell growth, glycerol consumption and ALS enzymatic activity

The effect of ALS inactivation on cell growth was examined during fermentation (Fig. 3A). The maximum OD600nm (biomass) reached was 6.80, and the growth of K. *pneumoniae* ΔALS was unaffected compared with the wild-type strain. Consumption of glycerol by K. *pneumoniae* ΔALS and K. *pneumoniae* HSL4 were also monitored in a fed-batch culture, and the glycerol consumption rate (5.50 g L^−1^ h^−1^) of K. *pneumoniae* ΔALS was slightly higher than that of K. *pneumoniae* HSL4 (5.34 g L^−1^ h^−1^), as shown in Fig. 3B. These results indicated that the consumption of glycerol was slightly promoted by inactivation of ALS, and further investigation revealed a notably lower ALS specific activity in the crude cell extract of *K. pneumoniae* ΔALS than the wild-type strain (Fig. 3C).

**Figure 3.**
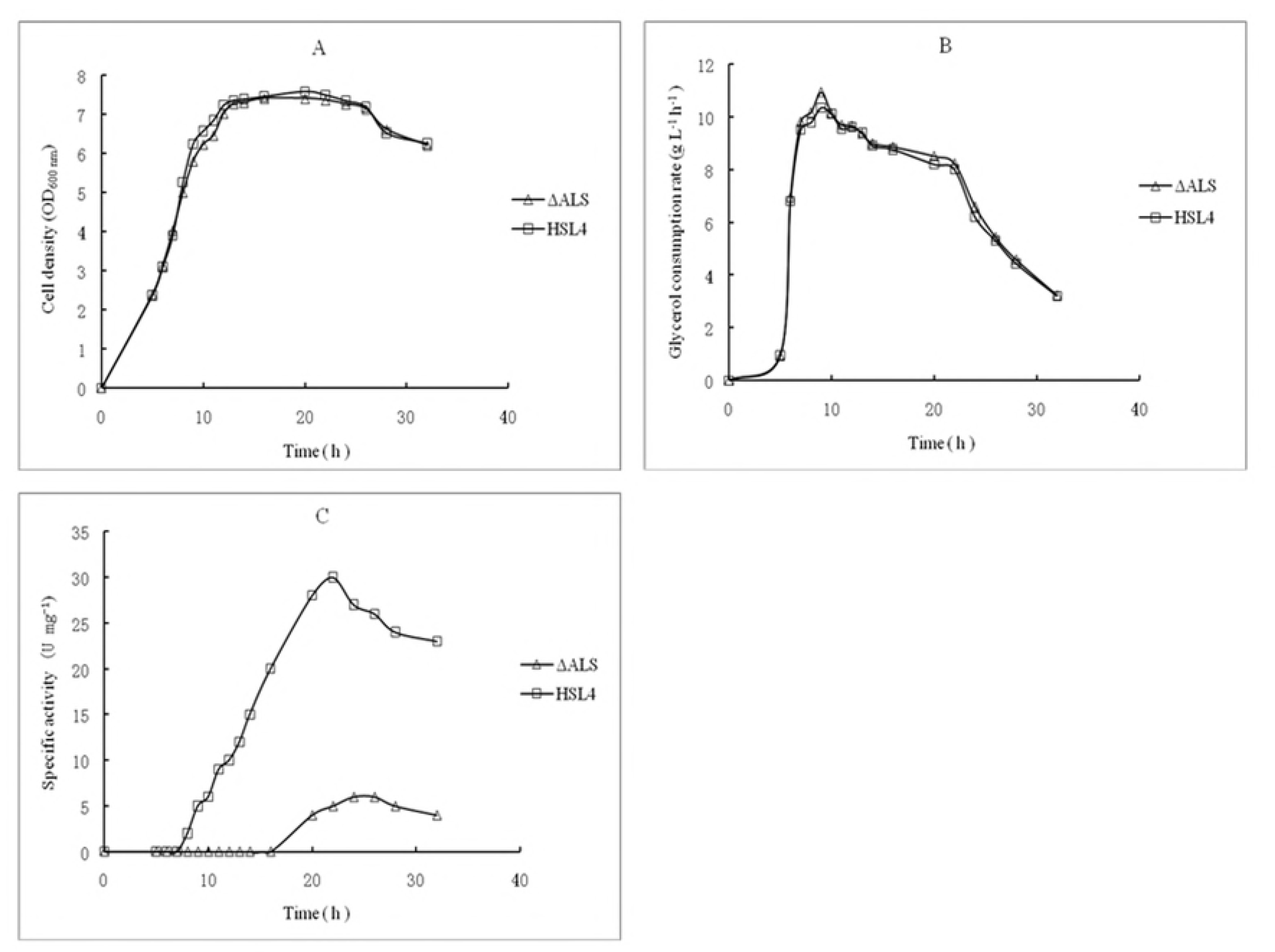
Time course of cell growth (OD600nm) (A), glycerol consumption rate (B), and ALS specific activity(C) of K. *pneumoniae* ΔALS and K. *pneumoniae* HSL4 in fed-batch culture. Enzyme activities are expressed in mM mg^−1^ min^−1^ (U mg^−1^).

### 3.3 Fed-batch fermentation of glycerol to 1,3-PDO by K. *pneumoniae* ΔALS

Fed-batch cultures were carried out in a 5 L fermenter to further investigate the fermentation of glycerol to 1,3-PDO by K. *pneumoniae* ΔALS, and the results after 32 h are shown in Table 3 and Fig. 4. After 5–16 h, the substrate was consumed fast and 1,3-PDO was rapidly synthesized by both K. *pneumoniae* ΔALS and the wild-type strain. However between 16–26 h, substrate uptake and the 1,3-PDO synthesis were slower, and both LAC and 2,3-BDO were rapidly accumulated by K. *pneumoniae* HSL4, while only LAC was rapidly accumulated by K. *pneumoniae* ΔALS. After 32 h, 1,3-PDO was the main product and reached a concentration of 72.04 g L^−1^ in the broth by K. *pneumoniae* ΔALS and 68.76 g L^−1^ in the broth of the wild-type strain. Fermentation by K. *pneumoniae* ΔALS produced five other main metabolites including 31.50 g L^−1^ LAC, 8.30 g L^−1^ SUC, 9.60 g L^−1^ AC, 7.60 g L^−1^ ETH, and 2.16 g L^−1^ 2,3-BDO. In comparison, K. *pneumoniae* HSL4 produced 15.42 g L^−1^ LAC, 7.76 g L^−1^ SUC, 7.02 g L^−1^ AC, 9.23 g L^−1^ ETH, and 13.75 g L^−1^ 2,3-BDO.

**Table 3.** Fed-batch fermentation of glycerol by K. *pneumoniae* ΔALS and K. *pneumoniae* HSL4 after 32 h.

| Strains | Substrate consumption |  | Metabolites (g L<sup>-1</sup>) |  |  |  |  | 1,3-PDO | 1,3-PDO |
| --- | --- | --- | --- | --- | --- | --- | --- | --- | --- |
|  | (g L<sup>-1</sup>) |  |  |  |  |  |  | productivity | conversion |
|  | Glycerol | 1,3-PDO | LAC | SUC | AC | BDO | ETH | (g L<sup>-1</sup> h<sup>-1</sup>) | (g g<sup>-1</sup>) |
| ΔALS | 175.91 ± 2.32 | 72.04 ± 1.46 | 31.50 ± 0.78 | 8.30 ± 0.28 | 9.60 ± 0.26 | 2.16 ± 0.15 | 7.60 ± 0.32 | 2.25 ± 0.03 | 0.410 ± 0.008 |
| HSL4 | 170.90 ± 2.24 | 68.76 ± 1.33 | 15.42 ± 0.89 | 7.76 ± 0.25 | 7.22 ± 0.21 | 13.75 ± 0.22 | 9.23 ± 0.19 | 2.15 ± 0.03 | 0.402 ± 0.007 |

Averages ± standard deviation were deduced from three experiments.

**Figure 4.**
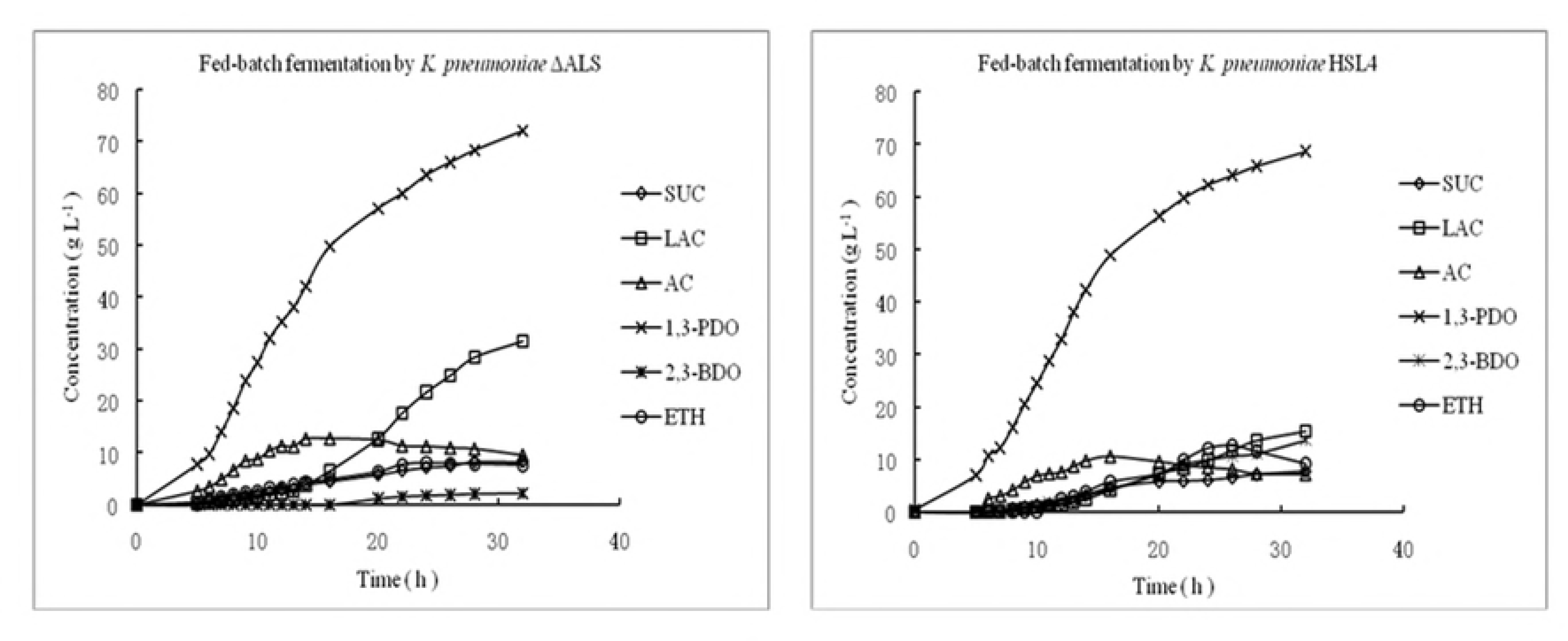
Time course of fed-batch fermentation of glycerol to 1,3-propanediol by K. *pneumoniae* ΔALS and K. *pneumoniae* HSL4 under micro-aerobic conditions. 1,3-PDO, 1,3-propanediol; LAC, Lactate; SUC, Succinate; ACE, Acetate; 2,3-BDO, 2,3-Butanediol; ETH, Ethanol.

As shown in Table 2, glycerol consumption by K. *pneumoniae* ΔALS was slightly higher than that of the parent strain. 1,3-PDO reached a final concentration of 72.04 g L^−1^ in *K. pneumoniae* ΔALS, which was an slightly increase of 4.71% compared to the parent strain. Similarly, 1,3-PDO productivity was 2.25 g L^−1^h^−1^ with K. *pneumoniae* ΔALS, higher than K. *pneumoniae* HSL4 (2.15 g L^−1^h^−1^). However, there was no notable difference in 1,3-PDO conversion between the strains. Therefore, as expected, ALS inactivation decreased 2,3-BDO generation, although K. *pneumoniae* ΔALS produced increased LAC (31.50 g L^−1^) than the parent strain (15.42 g L^−1^). This suggests that ALS deletion shifted the carbon flux away from the biosynthesis of 2,3-BDO towards the production of LAC.

### 3.4 Metabolic profiling of fed-batch fermentation by K. *pneumoniae* ΔALS

Based on the results of cell growth, substrate consumption and metabolite formation profiles (Figs. 3–4), fed-batch fermentations were divided into time four periods; (I) lag phase (0–5 h), (II) exponential growth phase (5–16 h), (III) stationary phase (16–26 h) and (IV) decline phase (26–32 h). Metabolic flux during different phases of the K. *pneumoniae* ΔALS and HSL4 fed-batch fermentations showed that during period I, cells were in a lag phase (Table 4). However, cells soon adapted to the new culture conditions, and the rate of substrate consumption and 1,3-PDO formation were high. Unfortunately, only a small amount of 1,3-PDO was synthesized due to the low biomass. During period II, cells grew exponentially, and both biomass and 1,3-PDO concentration increased sharply. Other metabolites were also synthesized steadily, but 1,3-PDO was the main product during this period. Acetate increased rapidly during the exponential growth period (5–16 h) and carbon flux was diverted to LAC synthesis in the stationary growth period (16–26 h), whereas 1,3-PDO synthesis slowed down during this phase. During periods III and IV, large amounts of LAC accumulated in K. *pneumoniae* ΔALS. Period IV is known as the decline phase, during which cell growth ceases, cells begin to die, and substrate uptake and metabolite formation decrease, although LAC synthesis continued in both K. *pneumoniae* ΔALS and K. *pneumoniae* HSL4. Other metabolites exhibited a similar flux distribution, although ethanol and acetate synthesis was negative due to volatilization and reutilization, respectively.

**Table 4.**
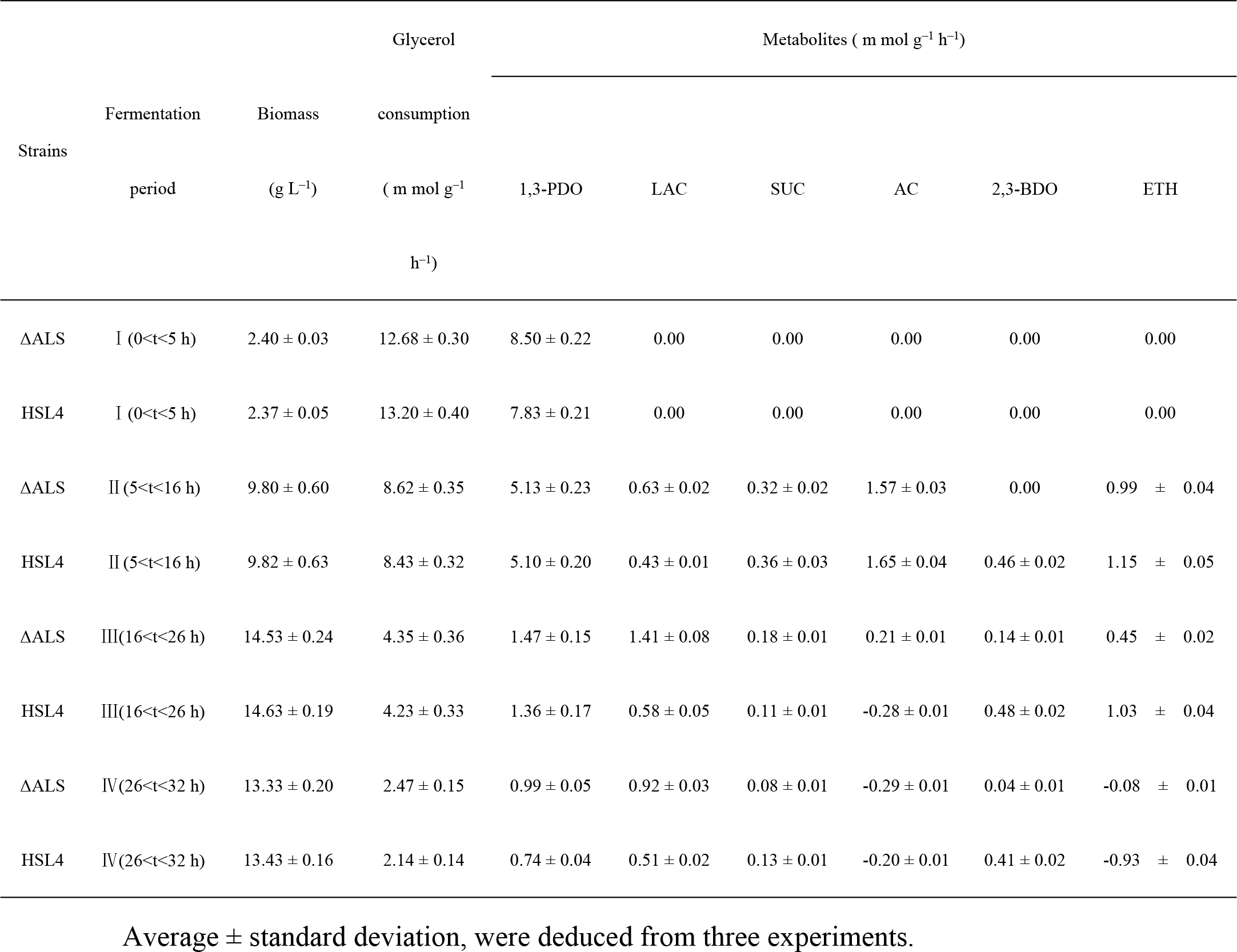
Metabolic flux in fed-batch fermentation by K. *pneumoniae* ΔALS and K. *pneumoniae* HSL4.

The distribution of metabolites in fed-batch fermentations by K. *pneumoniae* ΔALS and K. *pneumoniae* HSL4 (Table 4) clearly showed that the flow of glycerol to 1,3-PDO and byproducts other than 2,3-BDO and ETH was increased in the mutant K. *pneumoniae* ΔALS compared with the parent strain. It is worth noting that LAC increased strikingly in K. *pneumoniae* ΔALS, and it was reported that 2,3-BDO increased notably in a LAC-deficient mutant [12, 13]. In glycerol metabolism, LAC and 2,3-BDO reside on separate branches downstream from pyruvate and both depend on the action of dehydrogenase enzymes that use NADH as a cofactor. In general, one pathway is likely to be strengthened when the other is blocked, therefore it is logical that blocking 2,3-BDO biosynthesis resulted in the promotion of LAC biosynthesis in this study.

### 3.5 2,3-BDO stereoisomers in K. *pneumoniae* ΔALS and K. *pneumoniae* HSL4

*Klebsiella pneumoniae* synthesizes both *meso*-2,3-BDO and 2S,3S-BDO through fermentation of glycerol [17]. In this study, both mutant and the wild-type strains were cultured with glycerol (30 g L^−1^) as the carbon source, and 1,3-PDO was the main product with both strains (Fig. 5). However, the strains produced different amounts of 2,3-BDO of isomers; wild-type K. *pneumoniae* HSL4 produced 3.8 g L^−1^ *meso*-2,3-BDO and 0.20 g L^−1^ 2S,3S-BDO after fermentation for 12 h, whereas K.*pneumoniae* ΔALS produced 0.16 g L^−1^ and 0.85 g L^−1^, respectively. A distinguishing characteristic of the mutant strain was the low level of *meso*-2,3-BDO and high level of 2S,3S-BDO produced, suggesting synthesis of *meso*-2,3-BDO was inhibited while the formation of 2S,3S-BDO was stimulated in the absence of ALS.

**Figure 5.**
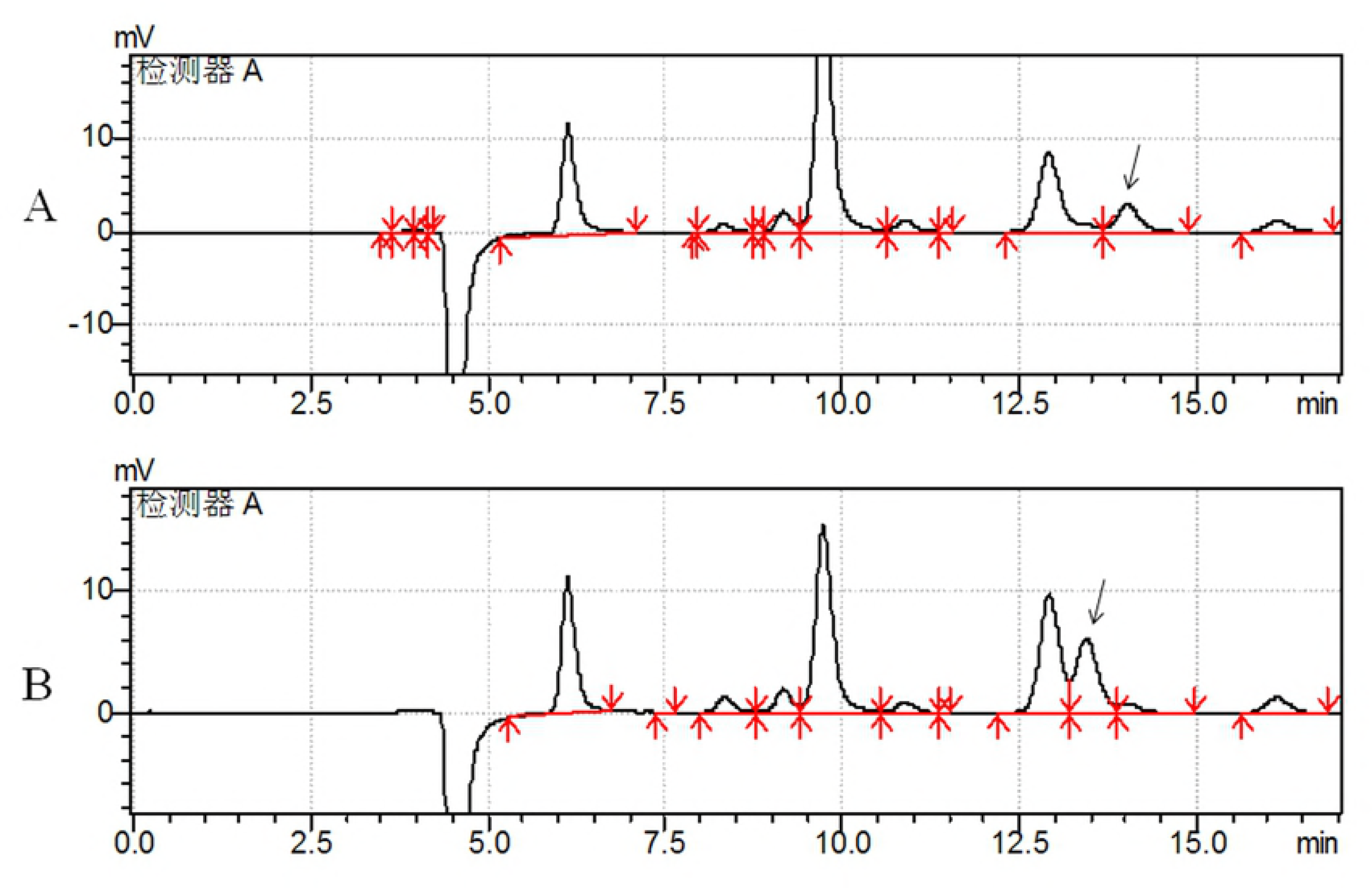
HPLC analysis of K. *pneumoniae* ΔALS (A) and K. *pneumoniae* HSL4 (B) fermentation broth following growth on glycerol. The arrows pointing to the peaks in (A) and (B) indicate 2S,3S-BDO and *meso*-2,3-BDO, respectively. Peaks corresponding to other metabolites and substrates are not indicated.

## 4 Discussion

During growth on glycerol in K. *pneumoniae*, considerable amounts of metabolites are produced via the activities of oxidative branch enzymes [10]. Lactate, 2,3-BDO, ETH, SUC and AC are among the major by-product metabolites generated, and these can account for 40%–45% of the total metabolites. In this study, LAC (15.42 g L^−1^) and 2,3-BDO (13.75 g L^−1^) were the two most abundant by-products and reached 22.4% and 20.0% (w/w) that of 1,3-PDO (68.76 g L^−1^), which were the major products in the fermentation broth of the wild-type strain. Minimizing the production of by-products is an effective way to engineer the microbial strain to synthesize more 1,3-PDO. Both ETH [11] and LAC pathways of K. *pneumoniae* [13] and *Klebsiella oxytoca* [12] have been successfully disrupted by knocking out genes encoding key enzymes in these pathways. Inactivation of the ETH pathway by knocking-out ALDH in K. *pneumoniae* YMU2 almost abolished ETH formation and significantly increased 1,3-PDO production [11]. Similarly, Yang et al. [12] reported that LAC-deficient mutants of K. *oxytoca* were obtained by knocking out the gene encoding LDH. Their results showed that 1,3-PDO concentration, productivity, and conversion rate can enhanced above levels achieved by wild-type strains, and LAC production can be diminished. However in their study, the distribution of metabolites were altered and over 60 g L^−1^ 2,3-BDO was produced as the major byproduct. Previously, the D-LAC pathway of K. *pneumoniae* HR526 was disrupted by knocking out D-LAC dehydrogenase [13], which increased conversion of 1,3-PDO and 2,3-BDO from 0.55 mol mol^−1^ to 0.65 mol mol^−1^, and the final LAC concentration decreased dramatically from more than 40 g L^−1^ to less than 3 g L^−1^.

To improve both the yield and productivity of 1,3-PDO from glycerol, disruption of the 2,3-BDO pathway is an alternative strategy. In K. *pneumoniae*, the genes coding acetolactate decarboxylase (ALD, encoded by the budA gene), acetolactate synthase (ALS, encoded by the budB gene) and butanediol dehydrogenase (BDH, encoded by the budC gene) catalyse the synthesis of 2,3-BDO [17]. The results of previous studies revealed that mutation of ALD not only decreased carbon flux to 2,3-BDO, but also increased the ratio of NADH to NAD+ by decreasing NADH consumption due to 2,3-BDO synthesis. Consequently, carbon flux was diverted towards the biosynthesis of 1,3-PDO [15]. Similarly, inactivation of BDH and expression of formate dehydrogenase reduced 2,3-BDO production and increased 1,3-PDO production [24], and disruption of the budC gene in K. *pneumoniae* ZG38 produced the same result [16]. Cui et al. [25] reported an improvement in the intracellular redox state in 2,3-BDO-deficient mutants lacking budA and B. Additionally, excess NADH was produced in the LAC-deficient mutant overexpressing dhaT. Since the synthesis of 2,3-BDO is catalysed procedurally by bud B, A and C and taking into account the reports that the mutation of budA and C promoted 1,3-PDO production, it is deductive that budB disruption should also be beneficial to the 1,3-PDO production. However, Oh et al. [26] reported the deletion of ALS gene (budB), hampering efficient 1,3-PDO production. The logical prediction was contradicted with the results by Oh et al. [26]. That implied the function of ALS gene (budB) refering to glycerol metabolism was deserved to be further investigated.

In this study, an ALS-deficient strain was constructed using the Red recombination system, and fermentation tests showed that inactivation of ALS caused a redistribution of metabolic flux, resulting in a decrease in the synthesis of 2,3-BDO and an increase in the synthesis of 1,3-PDO and LAC. Fermentation of K. *pneumoniae* ΔALS using glycerol as a substrate resulted in the production of 72.04 g L^−1^ of 1,3-PDO, which was a slightly higher than that of the parent strain (Table 3; Fig. 4). Inactivation of the 2,3-BDO pathway was speculatively associated with a decrease in the consumption of NADH under aerobic conditions (Fig. 1). This increases the available energy (NADH) supply that would otherwise be used by the competing 2,3-BDO pathway under anaerobic conditions. Moreover, LAC was the most abundant byproduct and reached 31.50 g L^−1^, double that of the parent strain. Cell growth was slightly decreased and glycerol utilization was slightly promoted compared to the parent strain. Interestingly, synthesis of *meso*-2,3-BDO was markedly inhibited in the mutant strain but synthesis of 2S,3S-BDO was increased (Fig. 5). This indicates that an unknown enzyme catalyzes the conversion of pyruvate to diacetyl. The mutant strain retained ~20% of the ALS activity of the parent strain, and this unexpected residual activity may have contributed to the results. Firstly, this may have complicated the enzyme assay which is based on measuring acetoin but not in the presence of acetolactate. Secondly, there may exist unknown (iso)enzymes that are functionally closely related to ALS, but this appears to be less likely.

Although the yield of 1,3-PDO was increased, the conversion was not promoted to a great degree, mainly due to the accompanying increase in LAC synthesis. Therefore, we predict that blocking LAC synthesis may be the preferred strategy when engineering 1,3-PDO production in *K. pneumoniae*. Our results indicate that 2,3-BDO synthesis competes with the synthesis of 1,3-PDO and LAC. Moreover, since blocking the formation of LAC stimulated 1,3-PDO and 2,3-BDO production in *Klebsiella* species [12, 13], we predict that knocking out ALS in combination with engineering of the LAC pathway would result in a mutant with lower LAC and 2,3-BDO by-products. This was proved by Oh et al. [26] but their did not get even higher 1,3-PDO production with the LAC-ALS dual-mutant. Heterologous expression of pyruvate decarboxylase and aldehyde dehydrogenase efficiently recovered glycerol metabolism in the 2,3-BDO synthesis-defective mutant [27]. They suggested the production of 1,3-PDO could be enhanced by preventing the accumulation of pyruvate.

Deleting the budB gene not only decreased the activity of ALS, but also decreased *meso*-2,3-BDO and increased 2S,3S-BDO. This phenotype indicates that *meso*-2,3-BDO formation is dependent on the activity of ALS whereas 2S,3S-BDO formation is inhibited by the activity of ALS. ALS is the first enzyme in the biosynthetic pathways of meso-2,3-BDO and 2S,3S-BDO in Enterobacteriaceae [17], and catalyzes the conversion of pyruvate to 2,3-BDO. Thus, it was not unexpected to observe a decrease in the formation of *meso*-2,3-BDO in the ALS-deficient mutant strain. The observed increase in 2S,3S-BDO in the mutant strain could be explained by an unknown enzyme that catalyzes the conversion of pyruvate to diacetyl and 2S,3S-BDO. Significantly, the mutation of ALS resulted in the accumulation of a greater amount of 2S,3S-BDO, suggesting this phenomenon could be helpful for 339 2S,3S-BDO production.

Glycerol metabolism is a typical biological redox pathway. In this study, a 2,3-BDO pathway-deficient mutant of K. *pneumoniae* HLS4 was constructed by knocking out the ALS gene. To investigate the influence of the ALS mutation on glycerol metabolism, ALS activity and various physiological properties including the yield of products and the distribution of metabolites and 2,3-BDO isomers was compared in the ΔALS and wild-type strains. Both the glycerol flux and distribution of metabolites were profoundly altered in the mutant strain in which the biosynthesis of *meso*-2,3-BDO was suppressed but 2S,3S-BDO synthesis was increased, while 1,3-PDO synthesis were slightly increased and LAC synthesis were largely increased. 1,3-PDO concentration and productivity were slightly increased by 4.71% and 4.65%, respectively. The overall conversion of glycerol to 1,3-PDO was not increased significantly due to an accompanying increase in the synthesis of LAC. Further engineering preventing the accumulation of pyruvate may be beneficial to improve 1,3-PDO production and reduce byproducts.

## Acknowledgements

This work was supported by grants from Guangdong Provincial Natural Science Foundation (2015A030313854), the National Natural Science Foundation of China (31502207) and Administration of Ocean and Fisheries of Guangdong Province (GD2012-D01-002).

